# The emergence of new lineages of the Mpox virus could affect the 2022 outbreak

**DOI:** 10.1101/2022.07.07.498743

**Authors:** Mayla Abrahim, Alexandro Guterres, Patrícia Cristina da Costa Neves, Ana Paula Dinis Ano Bom

## Abstract

Human Mpox is a highly contagious viral disease caused by Mpox virus, currently causing outbreaks in various regions of the world and considered a global public health problem. Understanding how the virus adapts or transfers to a new host species is crucial for effective prevention and treatment of viral infections. A comprehensive analysis of 1,318 accessible Mpox genomes was conducted to assess potential phenotypic effects, incorporating the examination of pseudogenes, gene duplication and deletion, as well as sequences obtained from the 2022 outbreak. This study we analyzed the genomic sequences of Mpox viruses from different clades and found that only a small percentage of proteins were conserved across the analyzed clades, with many proteins exhibiting various mutations, indicating the potential correlation between lineage or clade and variations in the pathology of human Mpox disease. The study highlights the importance of analyzing aligned regions in viral genomes to understand the evolution of the virus and the impact of structural changes on protein functions. In brief, further investigations are required to establish a correlation between the effects of these genetic modifications and the novel transmission pathways, average age, clinical manifestations, and symptoms observed in the ongoing outbreak.

## Introduction

In May 2022, numerous cases of Mpox started to be identified in several non-endemic countries. As of the previous week, at least 86,200 confirmed cases of Mpox have been reported across at least 90 non-African countries, resulting in 105 fatalities ^1–3^. *Mpox virus* is a double-stranded DNA virus with about 200-kb genome, being a member of the *Orthopoxvirus* genus from the *Poxviridae* family. These features are totally new for this disease in humans, since *Mpox virus* was endemic in West and Central Africa, and only occasionally caused short outbreaks elsewhere in the world, which were quickly contained or petered out by themselves ^4, 5^. In endemic African countries, published mortality rates vary from 1% to 10%. Despite the data restriction, the lineage/clade responsible for outbreaks in the Congo Basin appears to be associated with higher virulence ^6^.

In 2005, Likos and collaborators compared clinical, laboratory and epidemiological features of confirmed human Mpox case-patients. They suggested that human disease pathogenicity was associated with the viral lineage/clade (West African and Congo Basin (Central African)). A comparison of proteins between *Mpox virus* clades permitted the prediction of viral proteins that could cause the observed differences in human pathogenicity ^6^. Recently, two lineages of the *Mpox virus* were identified in the current outbreak in non-endemic countries. The most sequenced lineage/clade, to date, is related with a 2021 travel-associated case from Nigeria to Maryland in the USA (USA_2021_MD) that displays high similarity to the predominant 2022 *Mpox virus* outbreak sequences. The second lineage/clade is related to *Mpox virus* from a 2021 traveler from Nigeria to Texas in the USA (USA_2021_TX) ^7^.

The re-emergence and dissemination of the *Mpox virus* have resulted in infections across the globe. Something has changed. Before the 2022 outbreak, cases outside Africa have previously been limited to a handful that were associated with travel to Africa or with the importation of infected animals. Moreover, the ongoing cases differ from previous outbreaks in terms of age (thirties), sex/gender (most cases being males), and transmission route, being sexual transmission being highly likely. The clinical presentation is atypical and unusual, being characterized by anogenital lesions and rashes that relatively spare the face and extremities ^8, 9^. A comprehensive analysis was carried out on a total of 1,318 accessible Mpox genomes to assess potential phenotypic effects. The genetic analysis of the Mpox genomes is presented, incorporating the examination of pseudogenes, gene duplication and deletion, as well as sequences obtained from the 2022 outbreak.

## Methods

### Genome data, alignment, and annotation

Complete genome sequences of deposited *Mpox virus* were retrieved from the GenBank (www.ncbi.nlm.nih.gov/genbank) and GISAID (https://gisaid.org/) (accession 2023/02/03). A Perl script was utilized to carry out the following tasks: Firstly, the core reference genes and genomes were subjected to alignment using the BLASTn algorithm (https://ftp.ncbi.nlm.nih.gov/blast/executables/blast+/LATEST/). After alignment, structural annotation was carried out to determine the location of regions classified as putative Coding Sequence (CDS) or putative pseudogenes. The annotated regions were characterized according to the filters established in this study. A script was devised to identify the regions based on the following criteria: 1) the proportion of the aligned genome region to the size of the aligned reference gene; 2) percentage of identity; 3) alignment size relative to the reference gene size; 4) the proportion of the total aligned region divisible by three; 5) the presence of start and stop codons; 6) the presence of internal stop codons; 7) identification of nucleotide substitutions; 8) identification of nucleotide insertions; 9) identification of nucleotide deletions. Hence, the following filters were defined:

− **F1:** Alignment of genome region with 100% identity with reference gene.
− **F2:** Aligned genome region size equal to aligned reference gene size, with a percentage identity >= 80%, equal alignment size to reference gene size, total annotated region sizes divisible by three, and presence of nucleotide substitution.
− **F3:** Aligned genome region size not equal to aligned reference gene size, with a percentage identity >= 80%, presence of nucleotide substitution, and presence of nucleotide insertion.
− **F4:** Aligned genome region size not equal to aligned reference gene size, with a percentage identity >= 80%, presence of nucleotide substitution, and presence of nucleotide deletion.
− **F5:** Aligned genome region size not equal to aligned reference gene size, with a percentage identity >= 80%, total annotated region size not divisible by three, presence of nucleotide substitution, and presence of nucleotide insertion.
− **F6:** Aligned genome region size not equal to aligned reference gene size, with a percentage identity >= 80%, total annotated region size not divisible by three, presence of nucleotide substitution, and presence of nucleotide deletion.
− **F7:** Aligned genome region size not equal to aligned reference gene size, with a percentage identity >= 80%, presence of internal stop codon, total annotated region size not divisible by three, presence of nucleotide substitution, and presence of nucleotide insertion.
− **F8:** Aligned genome region size not equal to aligned reference gene size, with a percentage identity >= 80%, presence of internal stop codon, total annotated region size not divisible by three, presence of nucleotide substitution, and presence of nucleotide deletion.
− **F9:** Aligned genome region size not equal to aligned reference gene size, with a percentage identity >= 80%, total annotated region size not divisible by three, presence of internal stop codon, presence of nucleotide substitution, and presence of nucleotide insertion.
− **F10:** Aligned genome region size not equal to aligned reference gene size, with a percentage identity >= 80%, total annotated region size not divisible by three, presence of internal stop codon, presence of nucleotide substitution, and presence of nucleotide deletion.
− **F11**: Aligned genome region size equal to aligned reference gene size, with a percentage identity >= 80%, equal alignment size to reference gene size, total annotated region size not divisible by three, presence of internal stop codon, and presence of nucleotide substitution.
− **F12:** Aligned genome region size equal to aligned reference gene size, with 100% identity, and alignment size not equal to reference gene size.
− **F13:** Aligned genome region size equal to aligned reference gene size, with a percentage identity >= 80%, not equal alignment size to reference gene size, and presence of nucleotide substitution.
− **F14:** Aligned genome region size equal to aligned reference gene size, with a percentage identity >= 80%, not equal alignment size to reference gene size, total annotated region size not divisible by three, presence of internal stop codon, and presence of nucleotide substitution.
− **F15:** Alignments that do not fit the previous filters.

The regions that conformed to the filters F1, F2, F3, and F4 were designated as putative Protein Coding Sequences (CDS), while regions that met the filters F5, F6, F7, F8, F9, F10, F11, F12 and F14 were considered putative pseudogenes. The filters F13 and F15 may comprise a combination of coding DNA sequences and pseudogenes (Figure 01). These putative pseudogene sequences were characterized by the presence of internal stop codons and/or frameshifts. Subsequently, functional annotation of the putative CDS was performed by assigning the function of the most similar protein aligned in the region of interest. To validate the repeatability of the genome filtering method, a validation experiment was conducted utilizing the core reference gene set (CDS) against the reference genome nucleotide sequence.

**Figure 01.**
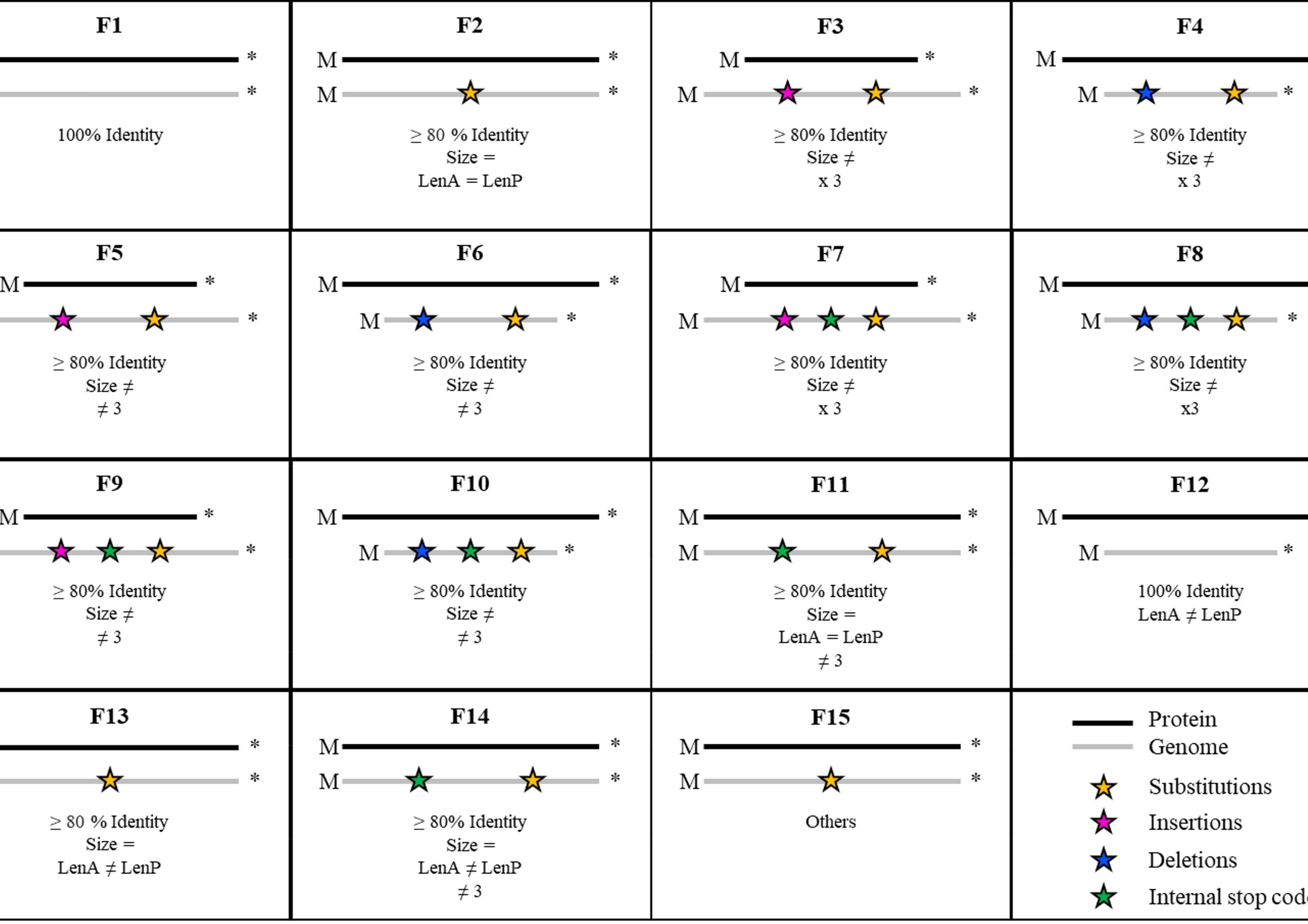
A schematic representation depicting the criteria utilized in each filter to ascertain CDS and putative pseudogenes.

## Results

A total of 3,272 *Mpox virus* genomes were acquired of GISAID and NCBI. After the download of the genomes, three filters were implemented to ensure improved reliability of the outcomes. These included: (1) Elimination of sequences labeled in the metadata as belonging to a certain clade or “probable” members of a different clade from the genomes acquired from GISAID; (2) Exclusion of genomes that had more than 10 instances of nucleotides marked as “N” through the second filter; and (3) Implementation of a maximum size filter to eliminate genomes with a genome size of over 200,000 nucleotides. During the application of the first filter, a sum of 113 genomes obtained from GISAD were eliminated. For the second filter, a total of 1,829 genomic sequences obtained from GISAD, in addition to 16 sequences from NCBI, were excluded. In the third filter, a total of 59 genomes from GISAD and 3 genomic sequences from NCBI were eliminated. As a result of these filtering steps, the final genomic sequence bank consisted of 1,318 sequences, out of which 1,268 were obtained from the GISAID database and the remaining 50 were from NCBI.

These genomic sequences have been assigned to 7 distinct clades based on their molecular characteristics. Among these, 40 genomes are assigned to Clade I, 10 genomes to Clade IIa, 01 genome to Clade IIbA.1.1, 07 genomes to Clade IIbA.1, 35 genomes to Clade IIbA.2.x (IIbA.2.1, IIbA.2.2, IIbA.2.3), 03 genomes to Clade IIbA.3, and 1,222 genomic sequences to Clade IIbB.1x (B.1.1 up to B.1.17). Clade I is equivalent to the previously designated “Congo Basin clade,” while Clades IIa and IIb correspond to the previously designated “West African clade.” The remaining clades comprise most of the genomes from human outbreaks that occurred in 2017, 2018, and 2022, with genomes from the 2022 outbreak falling within Clades A.2.x A3 and B.1.x^10^ (Figure 02). The set of core genes for *Mpox virus* consisted of 191 genes initially, based on the reference genome (NC_003310.1) of the *Mpox virus* strain Zaire-96-I-16 obtained from NCBI database.

**Figure 02.**
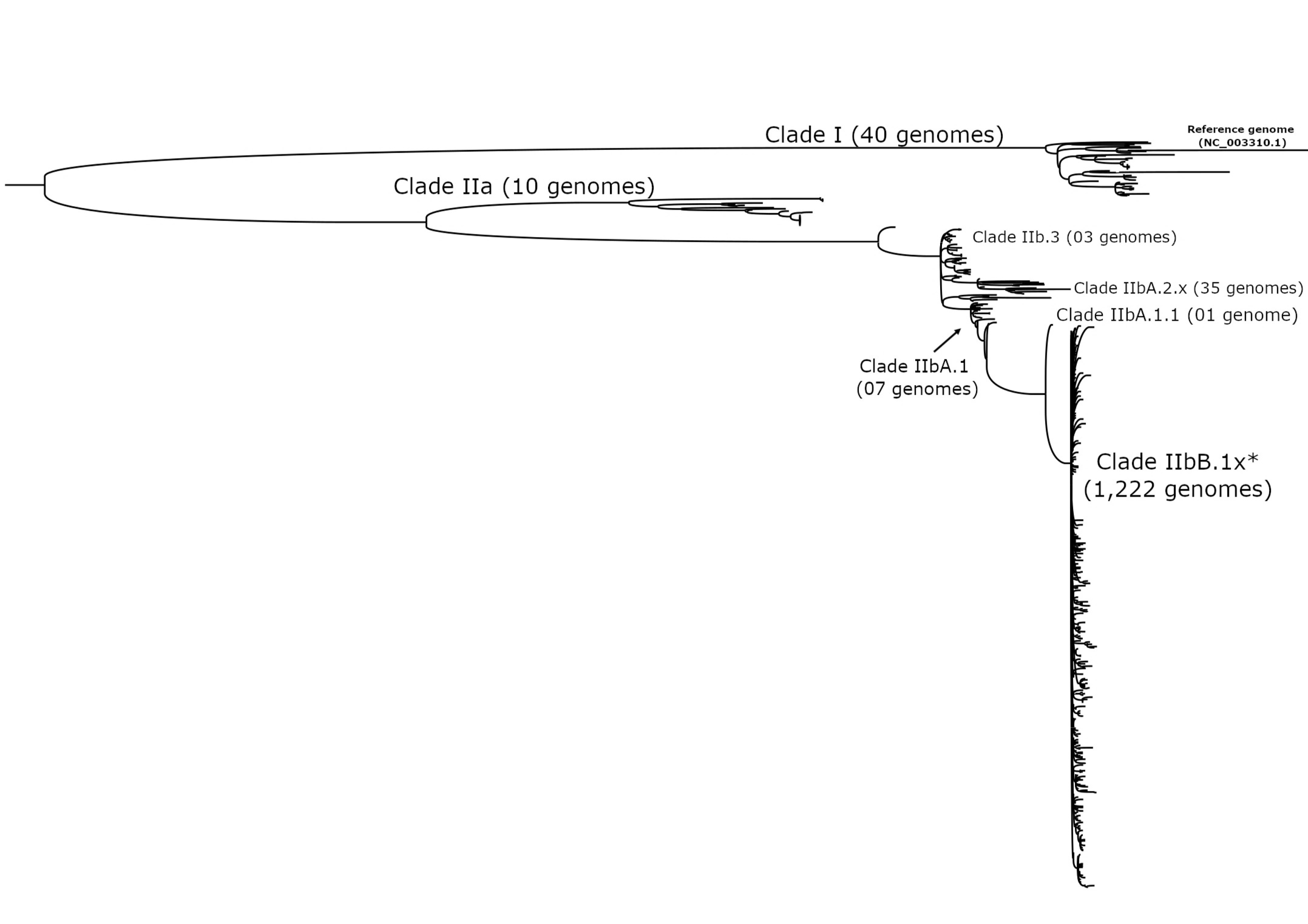
Phylogenetic relationships Mpox virus based on genomes complete sequences using the Maximum Likelihood Method using RAxML. Within the phylogenetic tree, we bring attention to the clades that were examined in our research and the total number of genomes that were included in each clade.

### Pipeline validation

By utilizing the file that contains the coding sequences (CDS) of the reference genome proteins, we conducted a search for the corresponding CDS in the reference genome nucleotide file. The search resulted in the identification of 191 proteins that exhibit 100% sequence identity between the query and the subject. These proteins were classified as belonging to Group F1, thus demonstrating the efficiency of the established pipeline in assigning accurate classifications.

Eight protein sequences, namely D1L (437 aa), J1L (246 aa), J1R (587 aa), J2L (348 aa), J2R (348 aa), J3L (587 aa), J3R (246 aa), and N4R (437 aa), were found to have duplications in the reference genome. The duplicated sequences exhibited 100% identity and were aligned at distinct coordinates, with the alignment size equal to the size of the respective protein. These proteins were also aligned on both the sense and antisense strands of the genome. However, it appears that certain proteins were mis- annotated as they were in the same genomic region and had identical sequences, specifically J1R and J3L, D1L and N4R, J1L and J3R, and J2L and J2R. The N3R protein was identified in the reference genome with a 100% identity match from position 190,103 to 190,633, spanning a length of 531 nucleotides, which is congruent with the original size of the protein. Additionally, a smaller fragment of the protein, with a length of 153 nucleotides and 100% identity, was observed aligned on the opposite strand (antisense) from position 6,226 to 6,378.

### F1 – Conserved proteins

A total of 142 proteins, accounting for 73.82% of the 191 examined, were detected in at least one of the *Mpox virus* genomes. Of the 191 proteins analyzed, only 13 were observed in over 94.99% (1,252 out of 1,318) of the genomes analyzed (Table 01). However, only 12 proteins are present in all clades (A23R, A30L, A3L, A42R, C13L, C20L, E9R, F10L, G3R, I2L, M5R and P1L). The remaining proteins were present in fewer than 97 genomes, constituting less than 7.35% of the total genomes analyzed. The maximum number of conserved proteins was discovered in Clade I, with a quantity of 142 proteins per genome, which constitutes 73.82% of the total 191 proteins analyzed. Notably, a total of 116 proteins were exclusively identified in Clade I. The number of conserved proteins in the other clades ranged from 13 in Clade IIbA.1.1 to 23 in Clade II.

**Table 01.** Of the 191 genes analyzed, only 12 were found to be highly conserved in a minimum of 1,252 out of 1,318 Mpox genomes evaluated.

| Gene | Amino acids (aa) | General function(s) of protein |
| --- | --- | --- |
| A3L | 77 | Hypothetical protein |
| A23R | 187 | Four-way junction DNA binding/ DNA recombination |
| A30L | 146 | Integral component of membrane/ viral envelope |
| A42R | 133 | Actin binding |
| C13L | 74 | Hypothetical protein |
| C20L | 73 | Hypothetical protein |
| E9R | 213 | Hydrolase activity |
| F10L | 129 | Virion core protein/ virion component |
| G3R | 220 | Late transcription elongation factor |
| I2L | 73 | Integral component of membrane/ virion membrane |
| M5R | 128 | Integral component of membrane/ virion membrane |
| P1L | 117 | Inhibits apoptosis and NF-κB activation |

**Table 02.** Out of the 191 genes that were examined, we identified 65 proteins that have the potential for pseudogenization. Moreover, a collective of 12 proteins were detected as having been altered in over 1,278 genomes.

| Gene | Amino acids (aa) | Protein name | General function(s) of protein |
| --- | --- | --- | --- |
| A38R | 212 | IEV transmembrane phosphoprotein | Cellular component/ plays critical role for actin tail formation |
| A48R | 84 | uncharacterized protein | Hypothetical protein |
| A50R | 554 | DNA ligase | DNA replication/ DNA repair/ DNA recombination |
| B1R | 70 | uncharacterized protein | Hypothetical protein |
| B3R | 299 | Serine/threonine-protein kinase 1 | serine/threonine protein kinase essential for transcription of viral intermediate genes |
| B14R | 326 | IL-beta-binding protein | Modulation by virus of host process |
| B16R | 352 | Glycoprotein | IFN-alpha/beta-receptor-like secreted glycoprotein |
| C9L | 487 | Kelch-like protein | intracellular protein expressed in late infection stages; may be involved in remodelling and stabilization of the host cytoskeleton |
| D11L | 153 | uncharacterized protein | Hypothetical protein |
| D18L | 107 | uncharacterized protein | Hypothetical protein |
| D2L | 64 | uncharacterized protein | Hypothetical protein |
| N3R | 176 | uncharacterized protein | Hypothetical protein |

A comparative genomics analysis conducted on the Clades (IIbA.2.x IIbA.3 and IIbB.1.x) implicated in the ongoing Mpox outbreak demonstrated the existence of a common set of 12 proteins in both clades. The occurrence of two distinct proteins (D6L and C12L) is restricted to a negligible frequency (<1%) in the genomes of Clade IIbaB.1.x. Clade A.2.x is uniquely marked by the presence of A14L and C3L proteins. Similarly, Clade IIbA.3 show the presence of A45L protein, whereas Clades IIbA.2.x and IIbB.1x exhibit the presence of C12L protein.

### F2 - Proteins that have mutated (substitution of nucleotides)

The F2 group of proteins exhibited a range of identity percentages, spanning from 96.05% to 99.96%. Out of the 191 total proteins, 179 (93.71%) were identified in at least one genome. Further analysis revealed that 143 (74.86%) of these proteins were present in more than 1,224 (92.86%) of the 1,318 total genomes analyzed. The remaining proteins were found in fewer than 70 genomes, comprising less than 5.31% of the total number of genomes analyzed. Clades demonstrated a comparable number of proteins, with 137 (71.72% of the total of 191 proteins) in Clade I and 157 proteins (82.19% of the total of 191 proteins) in Clade IIbB.1x. The analysis of the F2 filter classified proteins revealed that 102 proteins (53.40% of the total of 191 proteins) were present in all Clades. In contrast, 37 proteins were absent only in Clades I and II. Additionally, 23 proteins (12.04% of the total of 191 proteins) were present in a single clade, while 38 proteins (19.89% of the total of 191 proteins) were present in all Clades except Clades I and II. Clade I exclusively contained 19 proteins, while three proteins (B8R, G7R, and I5L) were present in 100% of genomes, and four proteins (A42R, C20L, E9R, and F10L) were exclusively present in Clade IIbB.1x.

The duplication of eleven proteins belonging to the F2 group, namely D1L, J1L, J1R, J2L, J2R, J3L, J3R, N1R, N3R, N4R, and R1R, was observed. Among them, D1L, J1L, J3R, and N4R were duplicated in all clades. J2L, J2R, and J3L were duplicated only in Clade I, while N1R, N3R, and R1R were duplicated only in Clade II. The duplicated sequences exhibited the same alignment size as their respective protein size and displayed an equal percentage of identity between the two duplicates. However, they were in different coordinates of opposite tapes, namely Sense and Antisense. The examination of the ongoing Mpox outbreak demonstrated the existence of 140 shared proteins in both clades. Additionally, a total of 14 proteins, namely A14L, A23R, A30L, A33R, A3L, A42R, B16R, C20L, C3L, E9R, F10L, G3R, M5R, and P1L, were discovered as being exclusively present in Clade IIbB.1x. Furthermore, two proteins (C13L and L4R) were detected exclusively in Clade IIbB.1x and IIBA.2.x.

### F3 - Proteins that have mutated (substitution and in frame insertions)

In the F3 group, we observed that the aligned regions had a percentage of identity ranging from 96.29% to 99.90%. Among the 191 proteins analyzed, 145 (75.91%) were present in at least one of the genomes. Six proteins (3.14% of 191 proteins) were found in more than 1,276 genomes (96.81% of 1,318 genomes): A28L, B17R, H5R, J1R, J3L and K1R. The remaining proteins were present in less than 35 genomes, which represents <2.65% of all genomes.

Upon analysis of the proteins in the F3 group and their distribution among the eight clades, it was discovered that three proteins, A28L, B17R, and K1R, were present in all clades. Furthermore, 49 proteins (25.65% of the total 191 proteins) were found to be exclusive to a single Clade. Clades IIbA.2.x and IIb.B.1.x exhibited the highest concentration of proteins, with over 100 proteins in each clade, while the remaining clades contained less than 11 proteins. Additionally, two proteins (1.04% of the total 191 proteins) were present in all eight clades except IIbA.1.1, which had only one genome. Finally, it was noted that B17R was present in all genomes, representing 100% of the studied population.

In the F3 group, 7 proteins were identified as being duplicated, including D1L, J1L, J1R, J3L, J3R, N2R, and N4R. The duplicated N2R proteins were observed to have an alignment size identical to the original protein size and the same percentage of identity between the duplications. However, they were in different coordinates on opposite strands, i.e., sense and antisense. The duplication of the N2R protein was found exclusively in Clade II. In relation to the Clades of the current Mpox outbreak, it was found that six proteins, namely A28L, B17R, H5R, J1R, J3L, and K1R, are present in both Clades. Additionally, 88 proteins are present exclusively in Clade IIbB.1.X and IIbA.2.x, while 33 proteins are only present in Clade IIbB.1.x (namely A18L, A24R, A39R, B10R, B14R, B1R, B21R, C1L, C5L, C7L, C8L, D16L, D19L, D3R, D5R, D8L, E9R, F5R, F6R, H1L, H6R, H7R, I1L, I3L, I7L, J1L, J3R, L2R, L4R, M2R, N2R, O2L, and R1R). Furthermore, 13 proteins are exclusively present in Clade IIbA.2.x, including A40L, A41L, A42R, A46R, A47R, B4R, C1L, C23R, D8L, E9R, F6R, H7R, and L2R.

### F4 - Proteins that have mutated (substitution and in frame deletions)

In the F4 group, the aligned regions showed a variable percentage of identity ranging from 83.12% to 99.84%. Among the 191 proteins, 144 (75.39%) were identified in at least one of the genomes. Out of these, 11 (5.75%) proteins were present in more than 1,265 genomes (95.97% of 1,318 genomes), which included A32L, A33R, B18R, B21R, B7R, C11L, C12L, D13L, D4L, J2L, and J2R. The remaining proteins were present in less than 52 genomes, which represented <3.94% of all genomes. Regarding protein duplication in the F4 group, 8 proteins were identified as duplicated in all Clades: D1L, J1L, J1R, J2L, J2R, J3L, J3R, and N4R.

Through analysis of the F4 group proteins and their relationship with eight different Clades, it was found that 123 proteins are present only in one Clade. Among these, 118 proteins were observed only in Clade IIbB.1.x and in less than 1% of the genomes. The D4L protein was found to be present in all Clades, with 100% occurrence, except in Clade I, in which it was present in only 2.5% of the genomes. Further, 6 proteins (A32L, B18R, B21R, D13L, J2L, and J2R) were present in 100% of genomes across all Clades except in Clade I, while 4 proteins (A33R, C11L, C12L, and B7R) were present in all Clades except Clade II. The IIbB.1.x Clade had the highest number of proteins with 140 (73.29% of the total 191 proteins) proteins. In the current Mpox outbreak, 11 proteins are present in both Clades, namely A32L, A33R, B18R, B21R, B7R, C11L, C12L, D13L, D4L, J2L, and J2R. Additionally, 127 proteins are exclusively present in Clade IIbB.1.x, while 3 proteins are present in both Clades IIbA.2.x and IIbB.1.x, namely B14R, C16L, and E9R. The C16L protein, however, is found only in clade IIbA.2.x.

### F5, F10, F12 and F13

We have observed a lower number of alignments in filters F5, F10, 12, and 13. From filter 5, only two regions aligned to the B14R protein were identified in two genomes of clade IIbB.1.x. The protein aligned with an identity percentage of 99.32%, on the positive strand, and contained only one gap, aligning in identical coordinates of the genome. The size of the reference genome protein is 981 nucleotides, however, the size of the alignment encompassed only the first 589 nucleotides. The F5 group showed no alignment at the end of the protein. We also found the B14R protein aligned in the same genomes classified in filter 15, with the alignment starting from nucleotides 591 to 981 of the protein, 34 nucleotides away from the end of the first alignment. The sorted alignment in filter 15 had three gaps and a percent alignment identity of 97.71%. Overall, the results demonstrate that the B14R protein had an initial alignment classified in filter 5 starting at a methionine (start codon) and ending at a stop codon. The protein may have “gained” 34 nucleotides, but lost a nucleotide (gap), resulting in a final gain of 11 amino acids without any changes in the reading frame. Additionally, the protein missed a triplet of nucleotides and ended its alignment with a stop codon in the sorted alignment in the F15 filter. The findings ultimately indicate that the area in question can be deemed a potential pseudogene, as evidenced by the presence of an internal stop codon, which is typical of pseudogenes.

Filter 10, which pertains to proteins that have undergone substitution, internal stop codon, and frameshift by deletion, revealed two proteins, namely C22L and A29L, in the genome of clade IIbB.1.x that were not classified in any other filter. The alignment region of C22L protein has an identity percentage of 96.81% and is smaller than the original protein by 36 nucleotides, while preserving the start and stop codons. However, 4 gaps were identified in this region that resulted in a change in the reading phase, leading to the generation of internal stop codons, which are indicative of pseudogenes. Meanwhile, the alignment region of A29L protein, which has a percentage identity of 97.88%, was found to have the same size as the reference protein (329 nucleotides) and contained preserved start and stop codons. Nevertheless, this region exhibited characteristics of a pseudogene with 2 gaps, which altered the reading frame through frameshift and generated an internal stop codon.

In filter F12, only the N3R protein was identified. Upon aligning the protein with the genome, it was observed that the size of the aligned region was smaller (455 nucleotides) than the original size (531 nucleotides) in two Clade I genomes. The aligned regions of the F12 group had 100% identity and preserved start and stop codons, however, the size of the region was not divisible by 3, indicating a possible change in the reading frame. In both genomes on the opposite strand (-), a portion of the protein with 153 nucleotides was aligned with 100% identity and a region size divisible by 3 in filter 15. However, despite having the stop codon preserved, the protein classified in this filter lacked a start codon.

A total of 7 proteins were classified in filter 13, displaying percent identity ranging between 91.18% and 99.84%. The region aligned to the L6R protein was present in all genomes of all clades. Both the A43R and N2R protein-aligned regions were found in 100% of the genomes of all Clades, except in Clade I, and showed duplication in Clade II. In contrast, the A48R protein-aligned region was present only in 2.5% of the genomes of Clade I. The B21R protein was identified exclusively in the Clades IIbA.2.x and IIbB.1.x of the current outbreak. Lastly, the B5R and G4L proteins were exclusively present in Clade IIbB.1x, accounting for less than 1% of the genomes.

### F15

This filter revealed that the aligned regions demonstrated variable percentages of identity, ranging between 85.77% and 100%. Out of the 191 proteins analyzed, a total of 65 proteins, constituting 34.03%, were detected in one or more of the genomes examined. Specifically, A38R, A48R, A50R, B14R, B16R, B1R, B3R, C9L, D11L, D18L, D2L, and N3R proteins were identified in over 1,278 genomes, which corresponded to 96.96% of the 1,318 genomes assessed. On the other hand, the remaining proteins were present in less than 68 genomes, representing less than 5.15% of all genomes. Regions aligned with 8 proteins (A10L, A28L, A34L, A8L, B10R, B14R, B21R and C16L) were identified as duplicates in different Clades.

The aligned regions of A10L were detected in 9 Clade 2 genomes (90% of Clade II genomes) and one Clade IIbB.1.x genome, with more than 93% identity. The regions were aligned on the negative strand and comprised two regions of the duplication. The first region was aligned between coordinates 1 and 235 of the reference protein and the other between 226 and 303, containing a repetition of 9 nucleotides (3 amino acids). However, there was a difference in coordinates of 27 nucleotides in the aligned regions of the genome. The region started at a methionine and ended at a stop codon, indicating that the protein might have experienced a gain of 27 nucleotides (9 amino acids) while maintaining the frame, with no gaps. These observations suggest that the protein may have undergone structural changes while still retaining its functional integrity.

Aligned regions of A28L were identified in the genomes of Clade I (10% of all Clade genomes), Clade II (100% of all Clade genomes), Clade IIbA.2.x (28.57% of all clade genomes), and IIbB.1.x (3.6% of all Clade genomes). Both genomes displayed a percentage of identity varying between 97.68% and 99.65%, as anticipated, and were found on the negative strand, with gaps ranging from 0 to 4 nucleotides. The coordinates of the protein sequences in the first alignment initiated at position 1 and concluded at coordinate 1,563 in the other alignment, which corresponds to the final coordinate of the protein, thereby indicating the preservation of the start and stop codons. Nonetheless, the coordinates of the aligned regions in the genomes were superimposed, indicating a frameshift mutation, which characterizes the region as a pseudogene.

Regions aligned to the A34L protein were identified in the two genomes of Clade II and one genome of IIbB.1.x, with a duplication of the protein present in the latter. In the IIbB.1.x genome, the first aligned region of the protein had a loss of three nucleotides, and a gap of 18 nucleotides (6 amino acids) was observed between the coordinates of the aligned regions. In addition, an insertion of 43 nucleotides was noted in the genome coordinates, and a total of 15 gaps were present in both alignments. Consequently, the region exhibited pseudogene characteristics with loss of start codon, frameshift, and no presence of an internal stop codon. In Clade II, the A34L protein was identified without being duplicated, with a percentage of identity of 99.77% and no gaps. However, the region exhibited a loss of the first 3 nucleotides leading to the loss of the start codon, which suggests the presence of pseudogene characteristics.

Aligned regions to the A8L protein were identified in two genomes of Clade IIbB.1x. In the genome with the gene duplication, start and stop codons were preserved in different regions of the aligned protein, but there was a loss of 11 nucleotides that caused a frameshift, and an internal stop codon was also present. As such, the region was identified as a pseudogene. In the other genome without duplication, there were 37 gaps and a loss of the first 8 nucleotides, resulting in a change in the reading frame. Regions aligned to the B10R protein were detected in 5 genomes of Clade II and 1 genome of Clade IIbB.1x, all of which exhibited loss of a region spanning 482 nucleotides of the protein. This finding signifies the pseudogenization of the region in all genomes.

Aligned regions to the B14R protein were identified in unique and duplicated regions in genomes of almost all Clades except for Clades I and II. In genomes where the B14R protein-aligned region appears once, the region showed an identity percentage of 97.71% with three gaps and a reduction in protein size to 393 nucleotides, which aligns with its alignment size. However, this region has lost the start codons and does not start at a start codon, but the stop codon is preserved, thereby characterizing the region as a pseudogene. The duplicated regions in the genomes exhibited identity percentages varying from 95.71% to 99.33% with gaps ranging from 1 to 7. The reference protein’s start and stop codons were preserved with overlapping of the genome regions, indicating a loss of part of the protein, which may characterize pseudogenization of the region.

The B21R protein-aligned regions were detected in a genome belonging to Clade I with an identity percentage ranging from 99.74% to 99.81%. The start and stop codons were conserved, however, the alignment exhibited a phase shift in the reading frame by 16 nucleotides in the genome and 15 nucleotides in the protein. Consequently, the region exhibited pseudogene-like characteristics. The C16L protein-aligned regions were identified in two genomes of Clade IIbB.1.x, one with duplication and the other without. The genome lacking duplication exhibited 97.51% identity, lost the last 4 nucleotides, and had 12 gaps, indicating pseudogenization. In the genome with duplication, the region was situated on the negative strand in contrast to the reference protein. Despite the percentage of identity varying between 99.58% and 99.4% in the duplicated region, the start and stop codons were preserved. However, it is likely that an assembly issue occurred as the protein was aligned at both the beginning and the end of the genome without loss of nucleotides. No proteins were identified for the F6, F7, F8, F9, F11 and F14 filters.

### Proteins that are classified under multiple filters

A total of 12 proteins exhibited aligned regions classified in more than one filter. Among them, the B14R protein was identified in both the F5 and F15 filters in two genomes of Clade IIbB.1x. Both genomes are located on the positive strand, with consecutive coordinates differing by 34 nucleotides and exhibiting a percentage of identity of 99.32% (F5) and 97.71% (F15), respectively. The aligned regions contained between 1 (F5) and 3 (F15) gaps. The size of the aligned region in the F5 filter was 589 nucleotides, while that in the F15 filter was 393 nucleotides. The total size of the reference protein was obtained by adding the two aligned regions together, resulting in 981 nucleotides. The D1L protein was identified in a Clade I genome and classified in both the F1 and F2 filters. Similarly, the N4R protein in the same Clade I genome was also classified in both the F1 and F2 filters. The regions aligned with these proteins are on opposite strands, have different coordinates, and are of the same size as the reference protein (1,314 nucleotides), with between 0 (F1) and 1 (F2) gap and percentage identity between 100.00% (F1) and 99.92% (F2). In addition, these regions were also classified in the F2 and F3 filters in the IIbB.1x genomes, where they maintained the previously described patterns, although the percentage of identity values varied between 99.61% (F2) and 99.54% (F3).

The J1L and J3R proteins were identified in the F2 and F4 filters, respectively, in distinct genomes of Clade IIbB.1x, located on opposite strands at different coordinates, and featuring between 0 (F4) and 2 (F2) gaps. The protein size ranged from 743 (F2) to 741 nucleotides (F4), in relation to the reference protein size of 741 nucleotides. Additionally, the percentage of identity varied from 97.84% (F2) to 97.43% (F4). The J1R and J3L proteins were subjected to the F2 and F15 filters, respectively, in different genomes of Clade IIbB.1x, and were found in different coordinates of opposite strands. The sequences had no gap, and their size and percentage of identity varied, with sizes ranging from 1,764 (F2) to 1,509 nucleotides (F4) in relation to the original protein size of 741 nucleotides, and percentage of identity varying from 99.94% (F2) to 100% (F4). Notably, the sorted region in the F15 filter lacked a stop codon. The J2L and J2R proteins were identified in the F2 and F15 filters in various genomes of Clade IIbB.1x, situated on opposite strands and with gap numbers ranging from 21 (F4) to 46 (F15). The region sizes differed from the original protein size (1,047 nucleotides), varying from 1,068 (F4) to 1,089 nucleotides (F15). The percentage of identity ranged from 87.87% (F4) to 93.91% (F15). The regions found in the F15 filter had no stop codon.

The M2R and M3L proteins were classified in filters F1 and F15, and F2 and F15, respectively, without any gaps, and in consecutive coordinates. The aligned region of the M2R protein is located on the positive strand and has sizes ranging from 279 (F1) to 52 nucleotides (F15), with 100% identity percentage. The region classified in the F15 filter lost the start codon. On the other hand, the aligned region of the M3L protein is located on the negative strand, with sizes varying between 1053 (F2) and 43 nucleotides (F15). The percentage of identity varies between 99.80% (F2) and 97.67% (F15). The region classified in the F15 filter lost the stop codon. The R1R protein, with an original size of 318 nucleotides in the reference genome, was classified in the F2 and F4 filters of Clade II genomes, located on opposite strands, with different coordinates and sizes ranging from 318 (F2) to 321 nucleotides (F4). The protein sequences contain between 0 (F2) and 3 (F4) gaps, with the percentage of identity varying from 98.42% (F2) to 97.19% (F4). The N3R protein was identified in almost all genomes, except for 21, and was classified in the following filters: F1 and F15 (20 genomes), F2 and F15 (1,245 genomes), F3 and F15 (13 genomes), F4 and F15 (20 genomes), and F12 and F15 (2 genomes)

### Gene deletion

The presence of four specific proteins was exclusive to the genomes belonging to Clade I (D14L, D15L, D16L and D17L), while none of the applied filters detected these proteins in the genomes of the other Clades.

## Discussion

Genomic surveillance has been vital to the early detection of mutations, monitoring of virus evolution and evaluating the degree of similarities between circulating. Molecular clock analyses assumed an evolutionary rate of 5 x 10^-6 11^. Our genomic analysis revealed that out of all the proteins examined, only 12 (6.28%) were conserved across the analyzed clades. Many of the proteins exhibited some form of mutation, including substitution of nucleotides, insertions, deletions, or even entire gene deletions. Several studies investigating single genes or entire genomes have indicated a potential correlation between lineage or clade and variations in the pathology of human Mpox disease. These findings collectively suggest that alterations in a relatively small number of genes may contribute to the modifications in viral clearance and pathogenesis observed in human infections ^5, 6, 12^.

Desingu and collaborators conducted a study on the evolution of the Mpox virus (MPXV) based on available sequence data. They found that the virus acquires point mutations in multiple proteins over time, leading to its evolution and the ability to cause outbreaks, such as the multi-country outbreak in 2022. The viruses exported from Nigeria to other countries in 2018-2019 were found to be evolutionary ancestors to the IIbB.1.x MPXVs causing the 2022 outbreak. The study revealed common amino acid mutations in 10 proteins among the exported viruses, with the MPXV/United States/2021/MD virus evolving from these mutations and further acquiring amino acid mutations in 26 proteins, including the 10 common mutations. Analysis of codon usage and host adaptation indices showed that the genes with nucleotide mutations in the B.1 lineage exhibited codon usage bias and were favorable for human adaptation, suggesting that selection pressure played a major role in their evolution. From 2017 to 2022, the MPXV’s continuous evolution through point mutations in its genes has been observed according to the available sequence data^13^.

We have observed a noteworthy trend in some genomes of Clade IIbB.1.x, where there is a higher proportion of proteins passing through filter 4 as compared to the other Clades. These proteins in this Clade exhibit a pattern of nucleotide substitutions and deletions. Americo and collaborators discusses how observed genome sequence variations may account for differences in virulence and transmission of MPXV Clades, but few such predictions have been tested experimentally due to the need for suitable animal models and stringent biosafety conditions. The study shows that MPXV Clades I, IIa, and IIb.1 exhibit large differences in morbidity and virus replication in CAST mice. The virulence differences between Clades I and IIa were less pronounced by the IN route, but replication in the lungs and abdominal organs still followed the order of Clade I > IIa > IIb.1. The determination of which genes are responsible for the virulence differences of MPXV Clades remains to be determined. The article also notes that recent gene loss has been a major factor in the evolution of orthopoxviruses that can account for differences in their host range. Both gain and loss of gene function should be considered in assessing virulence^14^. Our study revealed that four proteins are exclusive to Clade I of the Mpox genomes, and were absent in other Clades, including IIa and IIbB.1, as they had undergone gene deletion.

Regions aligned to A10L, A28L, A34L, A8L, B10R, B14R, B21R, and C16L proteins were identified as duplicates in genomes belonging to different clades. The study revealed that A10L underwent structural changes. The aligned regions of A28L, B14R, and C16L proteins exhibited pseudogene-like characteristics due to frameshift mutations, loss of start codons, and loss of nucleotides leading to changes in the reading frame. The A34L protein aligned regions exhibited pseudogene-like characteristics with loss of start codon, frameshift, and absence of an internal stop codon. Aligned regions of B10R protein were found in all genomes exhibiting loss of a region of the protein leading to pseudogenization. Regions aligned to A8L protein were identified in two genomes with pseudogene-like characteristics due to the loss of nucleotides causing frameshifts and the presence of an internal stop codon. Chen and colleagues demonstrated that MPXV-ZAI-V79 from the Congo Basin exhibited higher virulence towards cynomolgus monkeys compared to the presumed West African MPXV-COP-58. To elucidate the underlying reasons for this virulence difference, the team sequenced the genomes of one human West African isolate and two presumed West African isolates, and subsequently compared these sequences to that of MPXV-ZAI-96- I-16 from the Congo Basin. Through this analysis, they identified several proteins including proteins B10R and B14R along with other identified proteins (D10L, D14L and B19R), which may act as virulence determinants^15^.

In this investigation, Zhan and collaborators investigated alleles of 156 MPXV coding genes were gathered from roughly 1,500 isolates, including those responsible for previous outbreaks. Using molecular evolution methods, the study showed that homologous recombination has a minimal impact on MPXV evolution. Despite the majority of MPXV genes being subjected to negative selection, 10 genes were found to have sites evolving under positive Darwinian selection, most of which encode proteins interacting with the host and involved in host range determination. Most of these positive amino acid substitutions emerged several decades ago, indicating long-standing selection pressure, and three new positive substitutions appeared in 2019 or 2022. Protein modeling suggests that the positive amino acid substitutions could affect protein function^16^.

Researchers identified specific mutation sites that can be used as identification markers for different clades/lineages of MPXV. They found 84 mutations specific to Clades II and A, one SNP specific to Clades IIb and A, and one Clade IIb-specific mutation, among others. Two IIbB.1-specific SNPs (G76478A, G169968A) were also identified, which have not been previously reported. The study employed a random forest classification model to identify a combination of mutations that can distinguish different sublineages of IIbB.1 from other sublineages. Although the combination involves many mutation sites, it provides a method for scientific virus tracing. Overall, the findings provide insights into the evolutionary trajectory of MPXV outbreak strains and identify novel marker mutations for future studies on the transmission of MPXV strains^17^.

Guan and colleagues describe a study that investigates the global discrete phylogeographic analysis of MPXV, which is an orthopoxvirus that causes human Mpox. The study finds a strong association between MPXV and geography, implying that the virus has a strong evolutionary potential to adapt to the local ecology when it distributes to different regions. Positive selection analysis identifies two positive codons in the E13L scaffold protein and EEV membrane glycoprotein. The study also compares IFNα/βBP, a specific virulence protein of orthopoxviruses, and finds that the Ig I region of MPXV and VACV is relatively conserved. The study proposes that the function of MPXV IFNα/βBP and GAGs binding sites is maintained, and the amino-terminal portion of the IFNα/βBP has a lesser capability to inhibit IFN release, which may contribute to MPXV’s weaker pathogenicity. The findings could contribute to an expansive understanding of MPXV immune regulation and facilitate the design of intervention strategies^18^.

Irrespective of the underlying selective pressures leading to these mutations, it is reasonable to hypothesize that several mutations may have an impact on viral fitness. It is possible that a mutation that enhances one aspect of viral activity may diminish another. Although most current outbreaks of Mpox virus have resulted in mild disease symptoms, it is known to cause severe illness in specific populations, including immunosuppressed individuals, young children, and pregnant women ^19^. Although the association between pregnant women and the effects of human Mpox virus infection is poorly understood, evidence suggests that viruses belonging to the *Orthopoxvirus* genus are associated with increased maternal and perinatal morbidity and mortality ^20–22^. A recent study describes a cohort study of people living with HIV and low CD4 cell counts who were diagnosed with the Mpox outbreak between May 11, 2022, and Jan 18, 2023. The study included 382 cases from 19 countries and found that severe complications were more common in people with CD4 cell counts of less than 100 cells per mm3, including necrotizing skin lesions, lung involvement, and secondary infections and sepsis. Overall, 107 (28%) of 382 were hospitalized, of whom 27 (25%) died, all of whom had CD4 counts of less than 200 cells per mm3. The study suggests that Mpox in the context of advanced immunosuppression appears to behave like an AIDS-defining condition, with a high prevalence of fulminant dermatological and systemic manifestations and death^23^.

## Conclusion

Comprehension of how virulence develops during viral adaptation or transfer to a new host species is crucial for effective viral infection prevention and treatment. A better understanding of the evolution of virulence, obtained from various data sources such as phylogenetics, epidemiology, and experimental assessments of virus virulence and fitness, could lead to the development of novel approaches to control and eliminate human Mpox. Therefore, with the emergence of new lineages/clades, the evaluation of novel Mpox variants should include an assessment of the following questions: What effect do these mutations have on transmissibility and spread, antigenicity, aspects of pathogenesis, or virulence? Our study involved analyzing numerous full-genome sequences of Mpox viruses (MPXVs) sourced from different clades, including those implicated in the current outbreak. Our findings revealed that only a limited proportion of proteins were conserved among the studied Clades. Additionally, a multitude of proteins exhibited several mutations, such as nucleotide substitutions, insertions, deletions, pseudogenization, and even complete gene deletions, hinting at a plausible link between lineage or Clade and variations in human Mpox pathology.

## Author Approvals

All authors critically reviewed the manuscript for intellectual content and approved it in its final version.

